# diffloop: a computational framework for identifying and analyzing differential DNA loops from sequencing data

**DOI:** 10.1101/087338

**Authors:** Caleb A. Lareau, Martin J. Aryee

## Abstract

The three-dimensional architecture of DNA within the nucleus is a key determinant of interactions between genes, regulatory elements, and transcriptional machinery. As a result, differences in loop structure are associated with differences in gene expression and cell state. Here, we introduce *diffloop*, an R/Bioconductor package for identifying differential DNA looping between samples. The package additionally provides a suite of functions for the quality control, statistical testing, annotation and visualization of DNA loops. We demonstrate this functionality by detecting differences in DNA loops between ENCODE ChIA-PET datasets and relate looping to differences in epigenetic state and gene expression.

## BACKGROUND

The organization of DNA within the nucleus into hierarchical three-dimensional (3D) structures plays a key role in regulating gene expression by determining the accessibility of genes to the transcriptional machinery as well as the proximity of genes to their distal regulatory elements. Differences in 3D architecture, such as the presence or absence of “loops” between specific enhancers and their target genes, are associated with transcriptional variation in both normal and disease states [1]. Intriguingly, several recent studies have implicated pathogenic alterations in genome topology with a diverse set of diseases including cancers and autoimmune diseases [2–4].

Recent experimental techniques that couple chromatin conformation capture (3C)[5] with high-throughput sequencing have made the genome-wide identification of 3D interactions feasible. One such technique, Chromatin Interaction Analysis by Paired-End Tag Sequencing (ChIAPET), uses chromatin immunoprecipitation to capture interactions mediated by specific structural, regulatory, or transcriptional machinery proteins and is especially suited for high-resolution identification of 3D interactions. In addition, the high-throughput chromosome conformation capture (Hi-C) assay, which yields a potentially complete map of DNA proximity, can also be used to identify DNA loops [6, 7].

In order to fully explore the role that 3D genome organization plays in determining normal and pathogenic cell states, statistical tools are needed to identify differences in topology in a similar manner to which differential expression analysis is applied to transcriptional data. Additionally, the systematic integration of biological prior knowledge, such as the location of active enhancer regions, into topology analyses can provide annotation and insight into the regulatory role of a loop. Examples of loop annotation categories include enhancer-promoter [1] and disrupted neighborhoods[4]. Moreover, a computational framework that facilitates integration of other genomic data (e.g. DNA variation, RNA transcription, DNA and histone modification, etc.) in 3D genome analyses will help resolve the functional implication of DNA looping and relate these topological features to phenotypic variation.

To address these needs, we have developed *diffloop*, an R/Bioconductor package that implements statistical testing for differential DNA looping between samples from ChIA-PET and other 3D assays. *diffloop* additionally provides functionality for quality control and visualization of differential DNA topological features and facilitates a platform for integrative analysis with other types of genomic data. To demonstrate the utility of *diffloop*, we compare differences in DNA topology between three cancer cell lines using RNA Polymerase II ChIA-PET data from the ENCODE project. Subsequent integrative genomic analyses demonstrate the utility of incorporating topology data in analysis of epigenetic variation and gene expression.

## RESULTS

*diffloop* is designed to integrate with other bioinformatics packages available in the R/Bioconductor environment [8]. Although *diffloop* can identify topological differences and provide biological annotation of DNA loops for arbitrarily complex experimental designs, we focus for simplicity in this paper on two-group pairwise comparisons. Raw ENCODE project ChIA-PET sequencing data was obtained for five samples representing the MCF-7 breast cancer cell line (N=2), the K562 leukemia cell line (N=2), and the HCT-116 colon cancer cell line (N=1) [9].

### Preprocessing

Assays such as ChIA-PET enrich for chromatin interactions where interacting loci, termed anchors, are bound by a protein of interest. Anchors linked by paired-end tags/reads (PETs) represent distal regions of DNA that co-localize in three-dimensional space. The preprocessed data input to *diffloop* consists of interaction counts between putatively interacting anchors, as shown in **Figure 1**. Raw data preprocessing pipelines that produce the standard bed paired-end (.bedpe) input data format are described in the **Methods**. *diffloop* collates these read counts and assembles a list of anchors and a counts matrix (see **Methods)**. In effect, the counts matrix provides the level of evidence supporting each putative loop (row) for each sample (column). Under this construct, familiar techniques such as differential testing and principal component analyses can be applied to the counts matrix.

**Figure 1:**
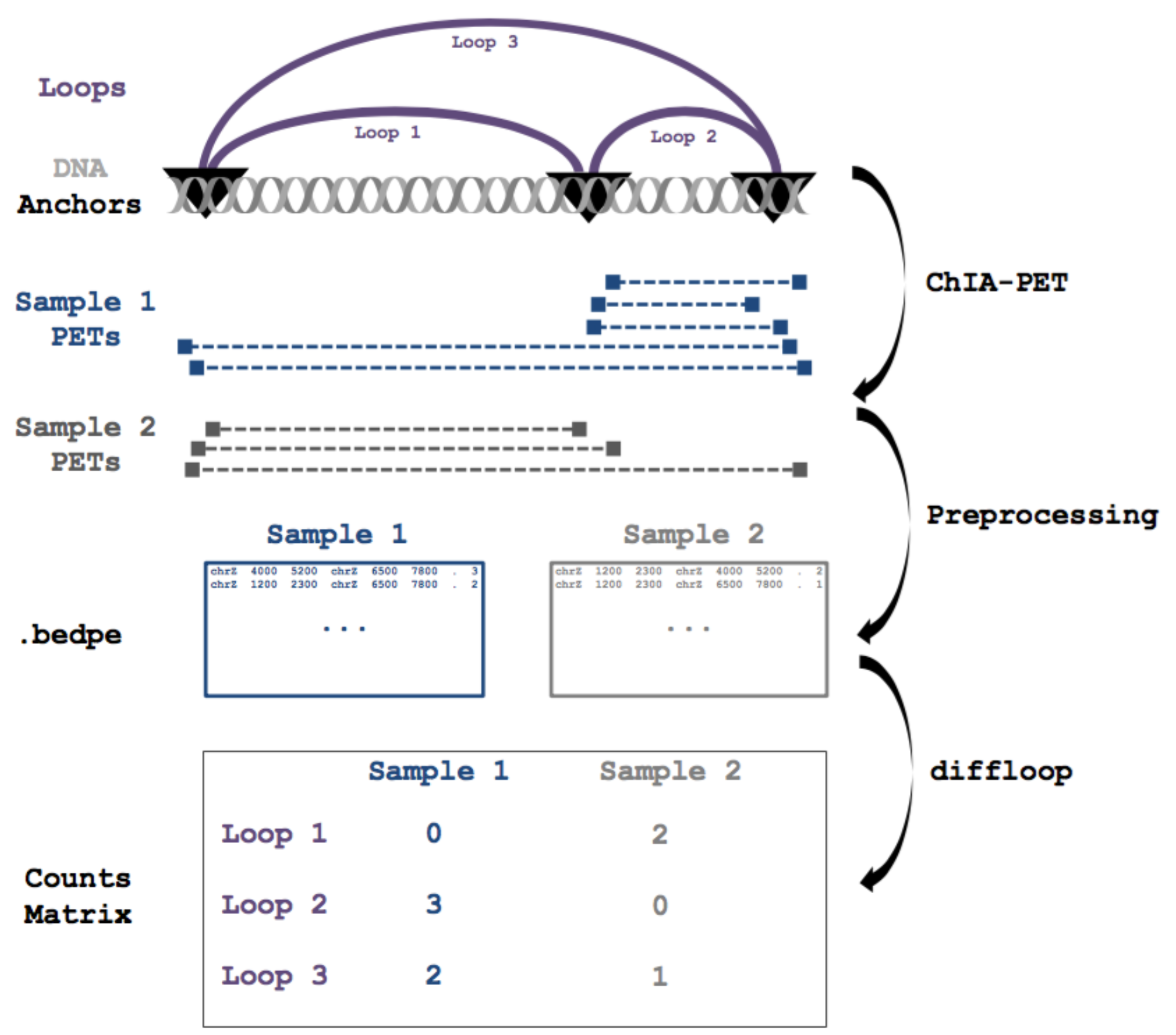
Overview of DNA looping data and its representation. The ChIA-PET assay generates sequencing reads that represent chromatin interactions associated with a protein of interest. Preprocessing involves identifying the interacting loci (loop anchors) and counting the number of reads supporting each interaction. These data are typically represented in .bedpe or a closely related format. The core functionality of *diffloop* imports these preprocessed loop data and assembles a counts matrix as part of a larger structure with metadata that can be used to assess differential looping.

### Quality Control and Normalization

Across all samples from the three cell lines considered for our analyses, we observed 89,806 anchor pairs involving 25,802 anchor loci. We first examined whether the cell types could be clustered using raw loop PET counts. To achieve this, we first computed the principal component plot of the loop counts matrix (see Methods). **Figure 2 (A)** reflects the first two principal components of the matrix with all of 89,806 loops. Counts are scaled by a per-sample size factor calculated using the DESeq2 normalization approach [10]. Scaling PET counts by these size factors removes the strong dependency of principal components on sequencing depth. A plot of the number of loops with varying levels of PET count support (**Figure 2 (B)**) indicates that the MCF-7 and K562 samples have similar coverage, while the HCT-116 sample appears as an outlier where very few loops are represented by multiple PETs. This lower quality sample was thus excluded from further analysis.

**Figure 2:**
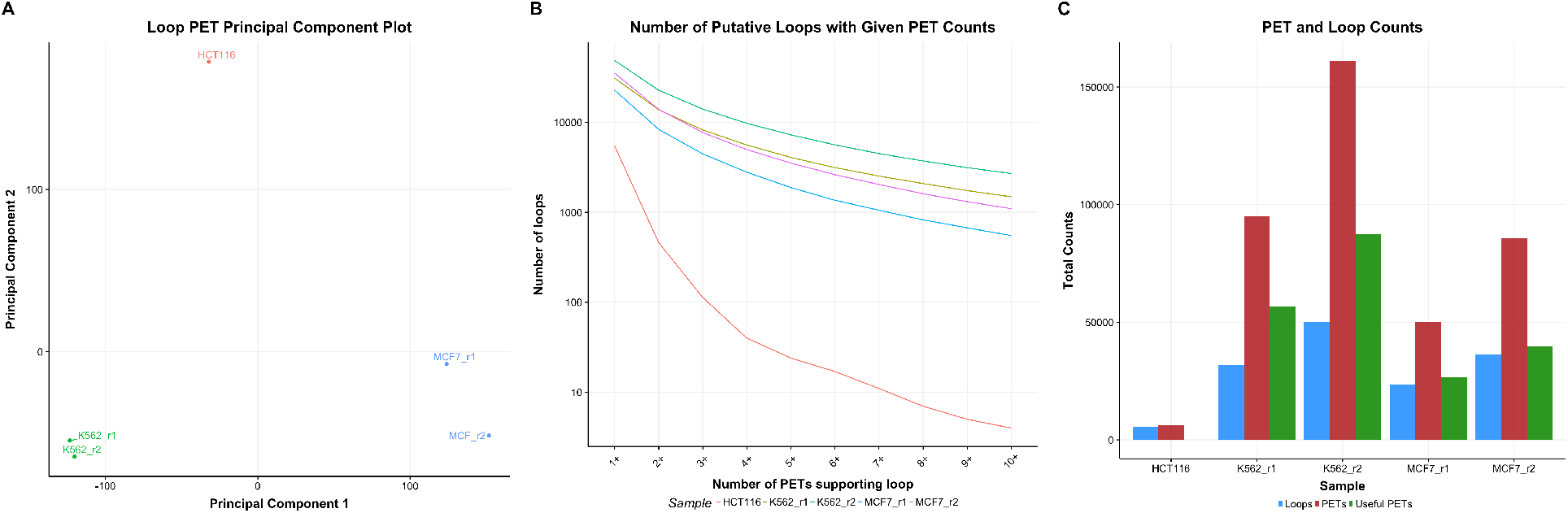
Principal component analysis and quality control summary. **(A)** A plot of the first two principal components of the loop PET count matrix. **(B)** The number of putative loops at varying levels of PET count support identifies the HCT-116 sample as an outlier. **(C)** The number of unique loops (blue), total PETs (red), and filtered PETs (green) supporting loops. The HCT-116 sample shows no filtered PETs as it was removed before quality control filtering on the loops.

As we wanted to exclude bias related to copy number variation (CNV) in these cancer genomes, we removed all interactions that were associated with anchors in known CNV regions in either of the MCF-7 or K562 cell lines. CNVs are known to affect chromatin interaction counts [11] and likely to bias differential looping analysis, especially in cancer genomes. In order to exclude such bias, we only considered interactions where neither anchor overlapped one of the nearly 500 CNV regions for either the MCF-7 or K562 cell lines. [9] Across all samples, we observed 9,723 of our 25,802 anchor regions overlapping a CNV region by at least 1 base and removed all loops involving at least one of these anchors, leading to 59,123 CNV-free loops in the K562 and MCF-7 samples.

We next filtered the observed interactions to retain only those that support valid loops suitable for differential analysis through a two-step process. First, we identified statistically significant loops by retaining only those anchor pairs whose interaction counts were significantly higher than that expected based on the background chromatin interaction frequency. *diffloop* combines counts across samples and assigns statistical significance to each putative loop using the method developed by Phanstiel *et al*. for the Mango ChIA-PET preprocessing pipeline [12]. We retained 24,601 loops in our combined K562 and MCF-7 samples that were significant at an FDR of 0.01 from the Mango model. Finally, we excluded anchor pairs not supported by at least two samples with two PETs each as proposed in other loop analyses [13], retaining 9,566 loops. Retaining loops with sufficiently high PET counts is analogous to filtering for genes that have a reasonable expression level in differential expression analyses [10]. **Figure 2(C)** summarizes the number of loops and PETs before and after these quality-filtering measures across each sample.

### Differential loop calling

To identify differential loops between cell types, *diffloop* applies the statistical test in edgeR [14] where counts are modeled using the negative binomial distribution and an empirical Bayes procedure is used to moderate the degree of overdispersion. The counts matrix, rather than representing reads mapped to genes or transcripts as is typical in an expression analysis, instead contains PETs (i.e. paired-end reads). We test for statistically significant differential loop strength (i.e PET count) between sample groups. While the mean-variance relationship of the ChIA-PET data does not deviate significantly from Poisson at current sequencing depths, the trend is similar to that observed at low transcript counts for RNA-Seq data (**Figure S1**). *diffloop* also makes the limma-voom [15] differential count test available as an alternative but we did not find evidence of improved performance relative to the default edgeR method (**Figure S6, S7; Table S3,S4**).

At an FDR of 1%, we identified 2,833 differential loops between the cell types, including 2,122 loops that were annotated as enhancer-promoter loops (see **Methods**). Of the 2,833 differential loops, 1,581 were more prominent in the MCF-7 (1,252 more prominent in K562). **Table 1** summarizes the top 5 differential enhancer-promoter loops unique to each cell line. To characterize the topological differences more systematically, we identified pathways enriched for genes involved in differential enhancer-promoter looping. We assessed MsigDB hallmark gene sets [16] using a Wilcoxon rank sum test (**Table 2)**. Several of the eight pathways enriched (FDR < 0.01) for genes with differential enhancer-promoter loops have existing evidence of relevance to the two cell types and cancers. Genes related to estrogen response such as GREB1 and XBP1 [17, 18], for example, are linked by several strong loops to unique enhancers in the MCF-7 breast cancer cell line. Conversely, targets associated with c-MYC transcription factor, which plays a well-documented role in leukemia [19], were enriched in K562. These results suggest that differential topology analyses can systematically uncover known and novel regulatory loops related to disease and other phenotypes of interest.

**Table 1:**
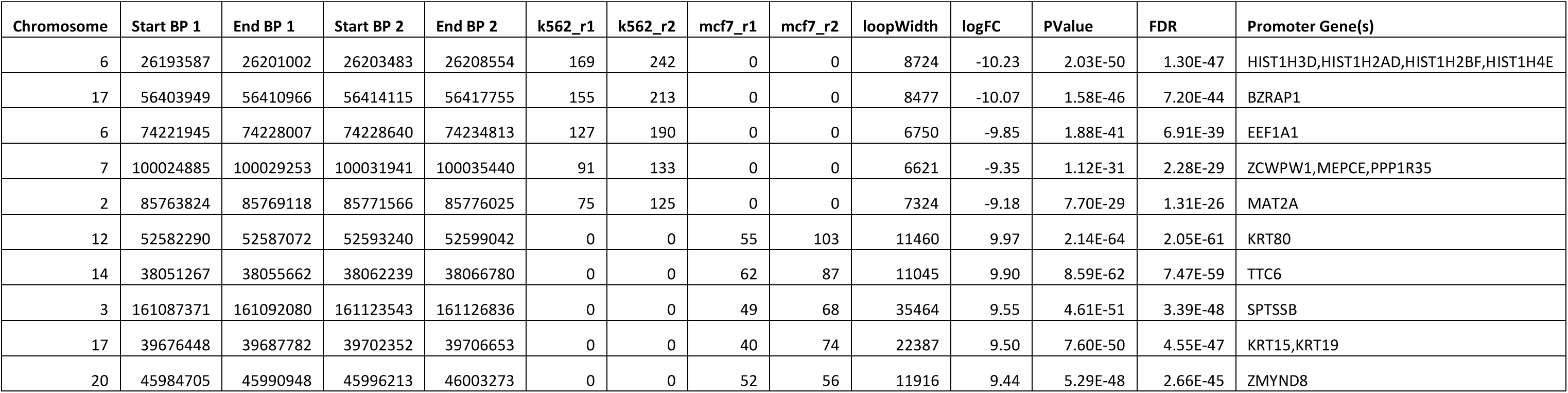
Top differential loops between MCF-7 and K562. The 5 most significant differential enhancer-promoter loops between these two cell types both more prevalent in MCF-7 and more prevalent in K562 are displayed. The loop annotation is listed alongside the summary statistics of each loop, which includes the number of reads that support each loop per sample. The final column lists all the genes of all promoter regions within 1kb of the loop anchor.

**Table 2:**
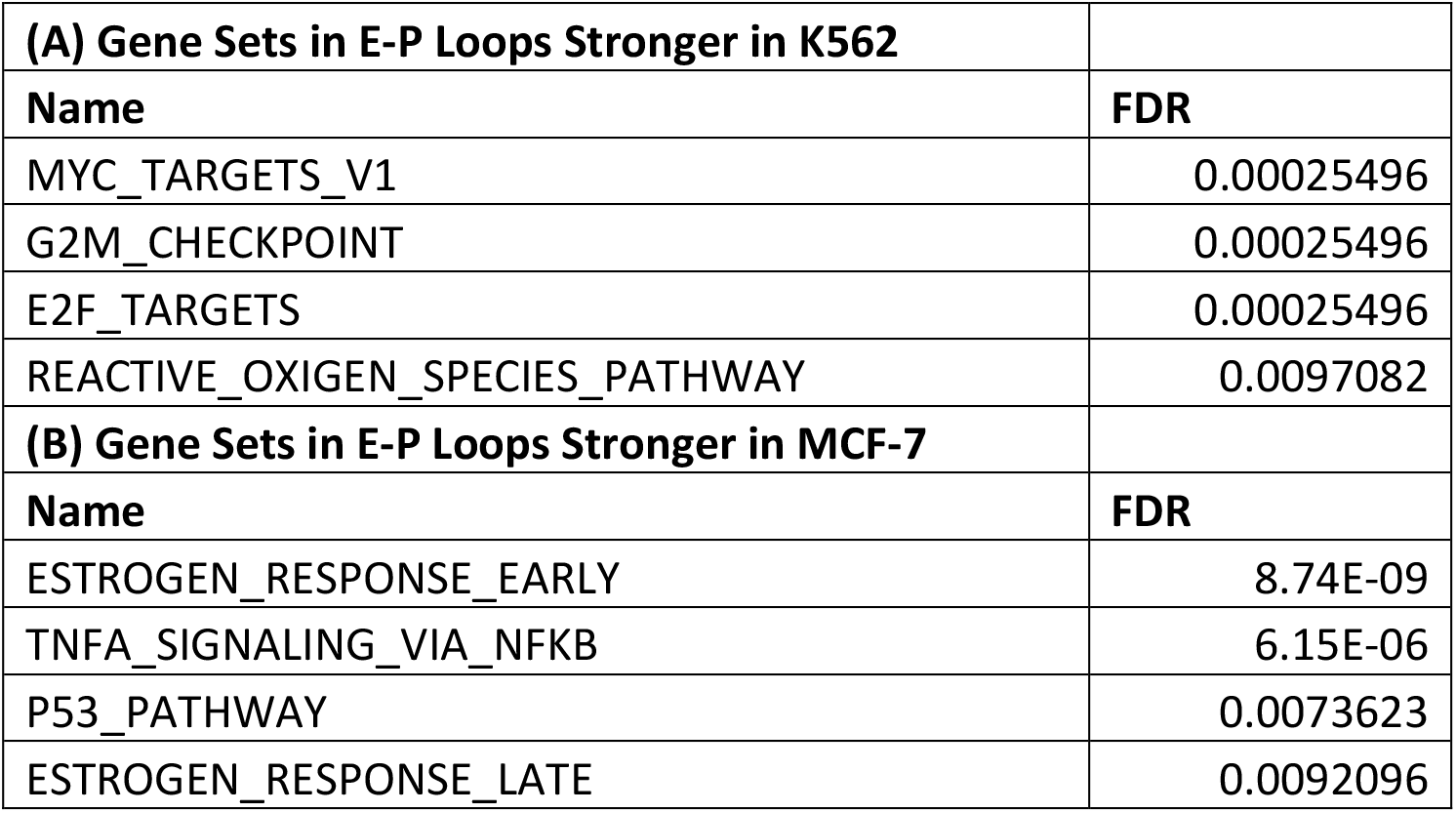
Pathway enrichment for differential enhancer-promoter loops. Querying the MsigDB hallmark gene sets [16] using a Wilcoxon rank sum test, we identified eight enriched pathways at an FDR of 0.01.

### Visualizations of differential looping

**Figure 3** shows two examples of regions with differential looping with corresponding tracks for the active enhancer mark, H3K27 acetylation (H3K27ac). Panel **(A)** shows a dynamic 3D landscape with many differential loops present more strongly in the K562 leukemia cell line than the MCF-7 breast cancer cell line. Most prominently, several unique enhancer-promoter loops (red) link the *MTHFR* promoter with nearby enhancers. Several of the enhancers are also linked by enhancer-enhancer loops (purple). Consistent with the increased looping the *MTHFR* gene is expressed at a higher level in the K562 (leukemia) cell line relative to MCF-7 (breast cancer) cell line (FDR = 2.95 × 10^−11^). Notably, variants near this gene have been associated with increased risk for a variety of leukemias.[20, 21] Conversely, Panel **(B)** shows a more active three-dimensional regulatory system in the MCF-7 cell line near the *NFKBIA* gene, which has been linked to breast cancer [22] and is overexpressed in MCF-7 compared to K562 (FDR = 0.0211).

**Figure 3:**
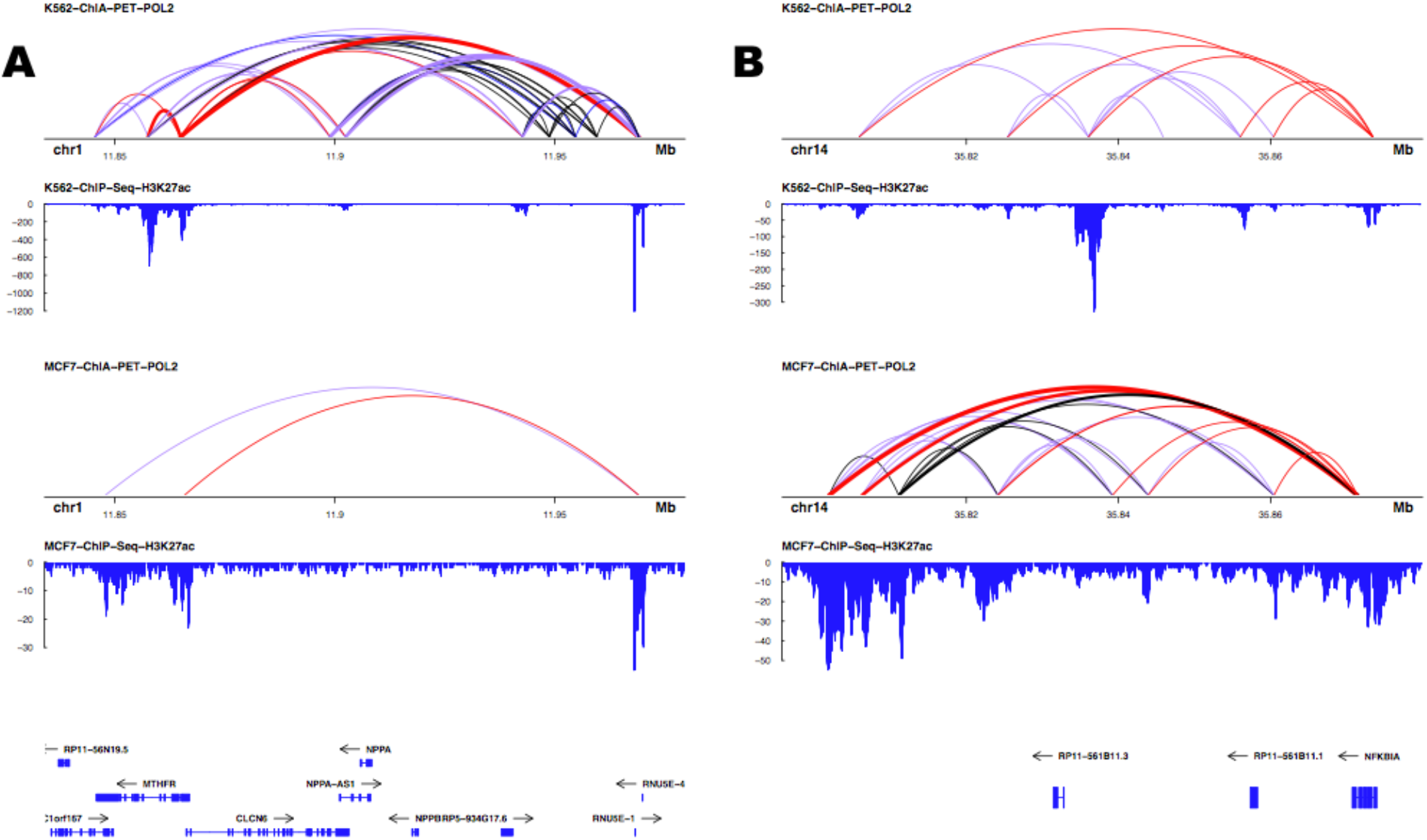
Sample visualizations of differential looping. Each panel shows the combined POL2 ChIA-PET replicates for the K562 and MCF-7 cell lines as well as the cell type-specific H3K27ac ChIP-Seq track. Line widths are indicative of the number of PETs supporting a loop while colors represent biological annotation (red: enhancer-promoter; purple: enhancer-enhancer; black: no special annotation). *MTHFR* (A) and *NFKBIA* (B) have variable topological and epigenetic landscapes between the two cell lines and have been implicated in leukemia and breast cancer, respectively.

### Epigenetic correlates of differential loops

To assess the relationship between differential looping and chromatin state, we compared the MCF-7/K562 log fold-change in PET count to corresponding ratios for DNase hypersensitivity (open chromatin), DNA methylation from the Illumina 450k array, and ChIP-Seq data for RAD21 (a cohesin subunit) and H3K27ac. As it has been suggested that the alteration of one loop anchor is sufficient to disrupt the loop[13], we used the anchor with the greatest difference in epigenetic mark in each case. For the sequencing data (ChIP-Seq, DNase), we subtracted the mean of the changes to account for variable total sequencing depth. **Figure 4** summarizes these results, which indicate that increasing loop strength in a cell type is positively correlated with **(A)** open chromatin, **(C)** cohesin localization, and **(D)** active enhancer markings. On the other hand, **(B)** DNA methylation was negatively correlated with loop strength, suggesting that hypermethylation of a genomic locus may inhibit loop formation.

**Figure 4:**
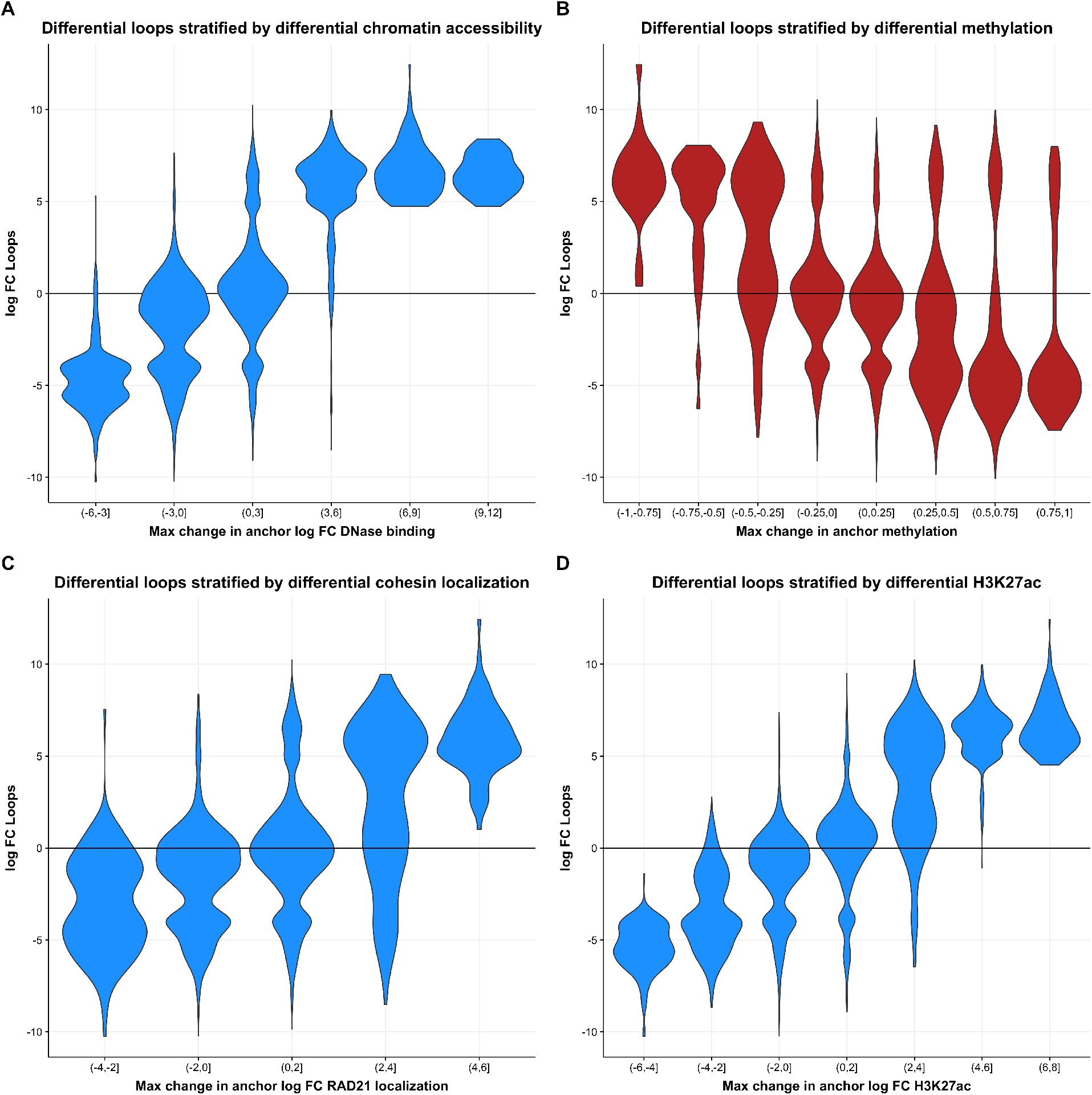
Differential loop strength correlated with various epigenetic factors. In each panel, the log2 fold change computed by *diffloop* for K562/MCF-7 is plotted against the log2 fold change of the epigenetic feature averaged over the region defined by the loop anchor. The anchor with the largest deviation from zero was chosen for each loop. Higher loop strength is associated with higher levels of **(A)** open chromatin, **(C)** cohesin binding, and **(D)** H3K27ac and lower levels of DNA methylation **(B)**.

### Relationship between looping and gene expression

Having ascertained that loop strength is tightly correlated with epigenetic state, we next sought to characterize the relationship between looping and transcription with a focus on the role of enhancer-promoter interactions. **Figure 5 (A)** summarizes differential expression in MCF-7 vs. K562 as a function of loop strength showing that differences in enhancer-promoter loop strength between the cell types is strongly positively correlated with differences in gene expression level. Of the pairs of enhancer-promoter loops/transcripts where both the loop and gene expression were significantly differential (FDR < 0.01; N = 144), 98.6% agreed in direction.

**Figure 5:**
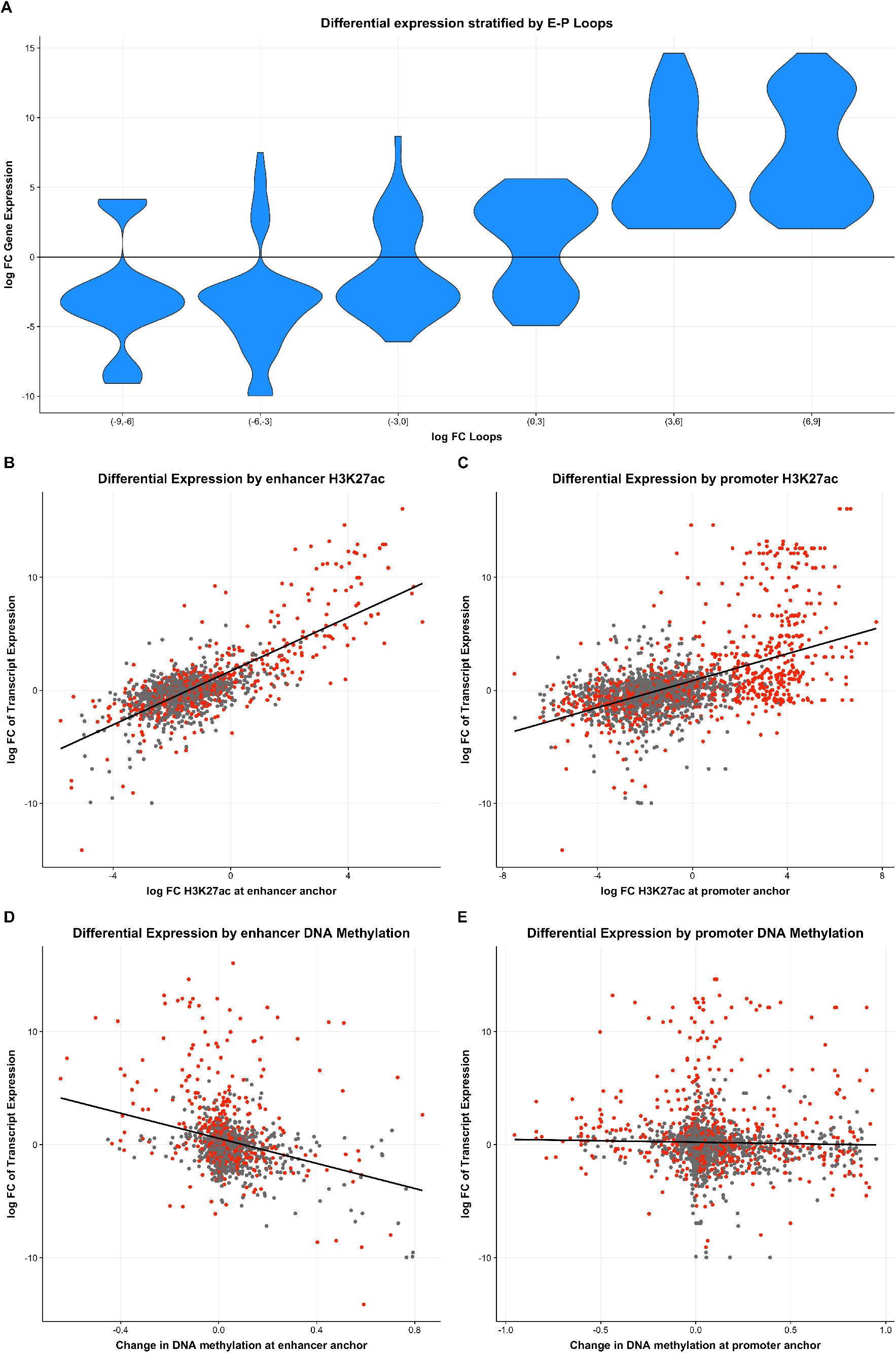
Effects of differential looping on transcription. **(A)** All loops that uniquely map to a single promoter region of differentially expressed transcript (FDR < 0.01). Increasing enhancer-promoter loop strength is positively correlated with increased transcription. We linked enhancer-promoter loops to their corresponding genes and assessed the effects of H3K27ac **(B)** distally and **(C)** proximally to the transcription start site. Similarly, the **(D)** distal and **(E)** proximal effects of DNA methylation were plotted against the expression fold-difference of the corresponding gene. Differential loops (FDR < 0.01) are highlighted in red in panels **(B)** - **(E)**.

As both H3K27 acetylation and DNA methylation are known to correlate with transcription[23], we sought to investigate whether these marks might play different roles at gene promoters vs. distal enhancers. **Figure 5** shows that, as expected, both distal **(B)** and promoter **(C)** changes in H3K27ac are positively correlated with expression whereas both distal **(D)** and promoter **(E)** changes in DNA methylation were negatively correlated with expression. Interestingly, the effect of epigenetic alterations was far more pronounced at the distal enhancer region compared to the promoter region. **Table 3** summarizes the simple linear regression and sample sizes for these panels as well as each plot contained in **Figure 5**. In particular, we note that variation in enhancer H3K27ac had the strongest correlation with gene expression. Given that identifying the enhancers associated with a gene is non-trivial, and that over 40% of enhancers skip the closest gene when interacting with their target promoters [24], this finding highlights the utility of a genome topology map for integrative analyses of transcription and epigenetics. Differential loops (indicated in red) did not appear to deviate from the overall trend of epigenetic values at enhancers/promoters correlated with the transcriptional outcomes.

**Table 3:**
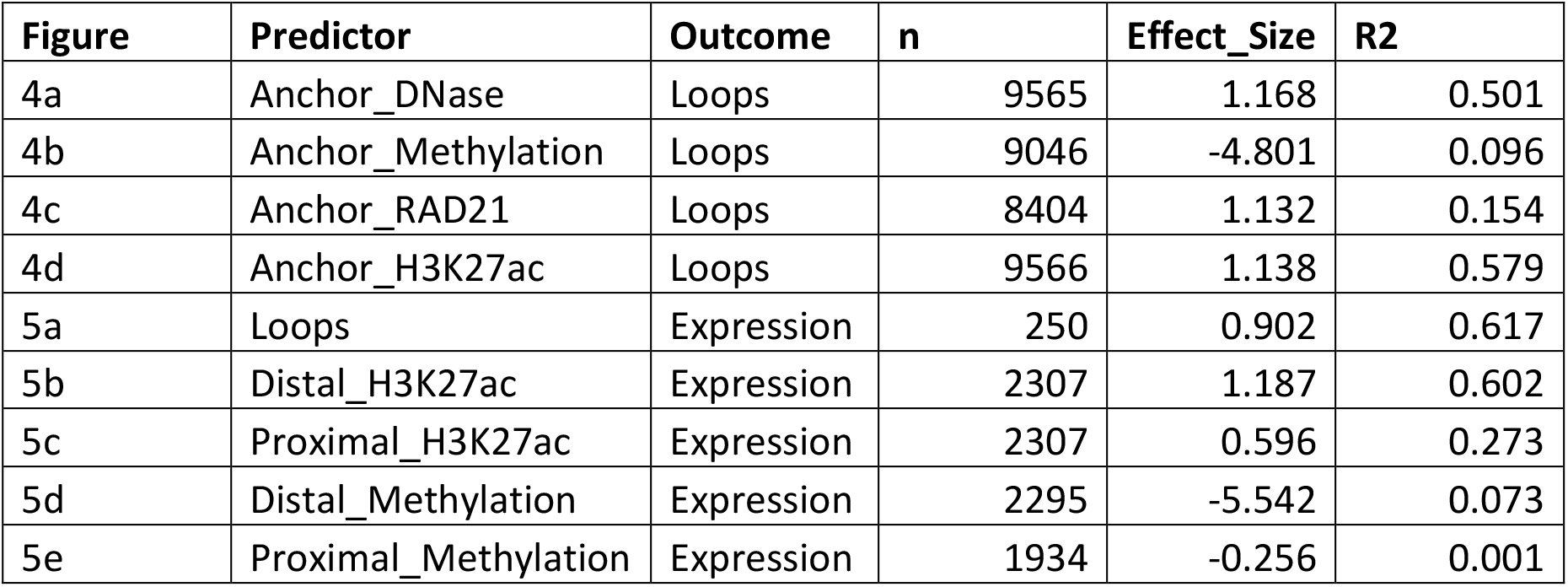
Model summaries of simple linear regression analyses of comparisons shown in **Figures 4–5**. The sample sizes, effect sizes, and *R*^2^ values of associations in **Figures 4–5** are listed alongside the Figure panel and variables used in the model. All models had a strong statistically significant linear term.

## DISCUSSION

A number of recent studies have implicated genome topology alterations in pathogenesis [1–4]. As the precise causes and effects of variability in DNA looping between cellular states is not well characterized, computational tools are needed to identify topological differences with statistical rigor and integrate putative regions of differential looping with the wealth of existing-omics data. To address this need, we have presented *diffloop*, an R/Bioconductor package that borrows much of its statistical methodology from differential expression analysis methods for RNA-Seq count data. Our package provides a full suite of functions to identify, annotate and contextualize DNA loops that vary between samples. The implementation of *diffloop* readily integrates with other R/Bioconductor packages and workflows and provides straightforward functionality for integrating genetic, epigenetic, and transcriptional data in the context of variability in the three-dimensional genome.

The base functionality of *diffloop* is designed to import processed data in a form that resembles .bedpe files (as shown in **Figure 1**) and perform differential loop calling two or more conditions that each have two or more samples. We note that integrating ChIA-PET data from experiments that target different factors is not advised. Moreover, while conditions without replicates can be analysed, *diffloop* will not assign statistical confidence for differences in this setting. Thus, we focused our analysis on ChIA-PET data from the K562 and MCF-7 cell lines that had replicates of ChIA-PET data against RNA POL2. Other analyses may focus on loops derived from ChIAPET protocols against other commonly studied factors such as CTCF, SMC1 or RAD21 as well as loops derived from Hi-C and similar chromosome conformation capture protocols.

We first compared the aggregate of the topological loops from ChIA-PET between three types of cancers and determined that cellular phenotypes indeed cluster based on their distinctive topology. However, as one of these cancer types (HCT-116) had only one technical replicate and had considerably poorer coverage we focused all subsequent analyses on the comparison of two cancer types with replicates, namely, K562 and MCF-7. While our comparison is close to minimal for statistical inference (2 groups by 2 replicates each), *diffloop* can handle arbitrarily complex designs, providing unique avenues to access the topological changes associated with cell lineages, such as differentiating stem cells [13] or pre-cancerous, oncogenic, and metastatic cell states.

Since chromatin interaction counts are known to be strongly distance dependent, we additionally investigated whether there could be between-sample variation in this relationship. This would suggest making the size factor dependent on loop width or loop PET count, similar to an approach used for Hi-C [25]. In this dataset, we did not find evidence for such a loop width dependency (**Figure S3**) and hence decided that a single scale factor correction value for each sample was sufficient for normalization of the examined datasets.

When assessing the correlation between variability in epigenetic features (open chromatin, DNA methylation, histone acetylation, and cohesin localization) and differential loops (**Figure 4**), we determined that strong changes in these one-dimensional epigenetic data at either anchor were highly correlated (**Table 3**) with the presence of a differential loop. It is possible that significant alterations in DNA methylation or open chromatin may affect the ability of loop-mediating proteins to bind to the genome, and that resulting loss of a DNA loop prevents the transcriptional machinery from activating target genes. We therefore expect that the topological structure of the genome is a vital link in understanding the effect of epigenetic variability on gene expression. Similarly, much of the effect of distal genetic variation on transcription as uncovered through expression quantitative trait loci analyses may be mediated through the 3D architecture of the genome.

One key finding was that variation in distal DNA methylation and distal enhancer markings (indicated by H3K27ac) are more highly correlated with differential expression than the epigenetic signature at proximal promoter regions. Our finding corroborates previous findings that enhancer DNA methylation more highly correlated with gene expression [26], and provides a framework to use topological data to conveniently link distal enhancer regions to their specific transcription start sites. While the distal regulatory elements accounted for a significant proportion of the variability in gene expression (**Table 3**), the complexities associated with multiple enhancer-promoter and enhancer-enhancer loops per transcription start site convolutes the direct effect of epigenetic variation on gene expression. We anticipate that subsequent iterations of *diffloop* will include functionality to synthesize the full spectrum of connections when linking epigenetic data to transcriptional variation as mediated through the 3D genome.

An estimated nearly 40% of enhancer regions affect genes that are not the closest (in one-dimension)[24] implying analysis of the topology of the genome is vital to understanding what epigenetic modifications affect transcriptional processes. Moreover, as significant variability in transcription leads to distinct cellular states, characterizing the variable topology of the genome is vital for mechanistically bridging epigenetic changes like variable open chromatin and DNA methylation to their transcriptional effects. While 3C assays have provided a means to localize distal regulatory regions in the topological landscape to specific genes, statistically rigorous and computationally flexible tools are needed to fully characterize the 3D genome. Our analyses of the topological variability between cancers from the ENCODE project suggest that *diffloop* may be a useful resource for integrating genome topology data into existing workflows.

## CONCLUSIONS

Our novel software package, *diffloop*, provides a user-friendly environment for analyzing genome topology data in the R/Bioconductor environment. Specifically, *diffloop* provides a suite of tools to uncover differential loops in DNA with statistical rigor and integrate other bioinformatics data. Our analyses show how differences in chromatin accessibility, DNA methylation, histone modifications, and cohesin localization correlate with differences in DNA looping, and how looping relates to differences in gene expression and cellular phenotypes. In particular, epigenetic variability in distal regulatory elements is more tightly correlated with gene expression than in promoter regions, emphasizing the value of an improved understanding of the 3-dimensional structure of the genome.

## METHODS

### diffloop Data Structure

The core functionality of *diffloop* is designed around *loops* objects, a novel S4 class implemented in the package. Each *loops* object contains five slots that provide an efficient storage of three-dimensional data in the R environment. Specifically, the anchors slot contains a GRanges[27] object specifying the genomic coordinates of the DNA anchors identified through peak calling; the interactions slot represents each loop (row) by a pair of indices (two columns) specifying its two anchors; the counts matrix summarizes the number of supporting PETs for each loop (row) per sample (column); the colData slot contains per-sample information, such as group labels and normalizing constants; and the rowData slot, which provides per-loop annotation, such as loop width, loop type, and statistical significance measures. **Figure 6** provides a graphical overview of the loops object as well as the functions (indicated in bold) employed in *diffloop* to integrate the heterogeneous data and the additional software packages that readily integrate with our framework.

**Figure 6:**
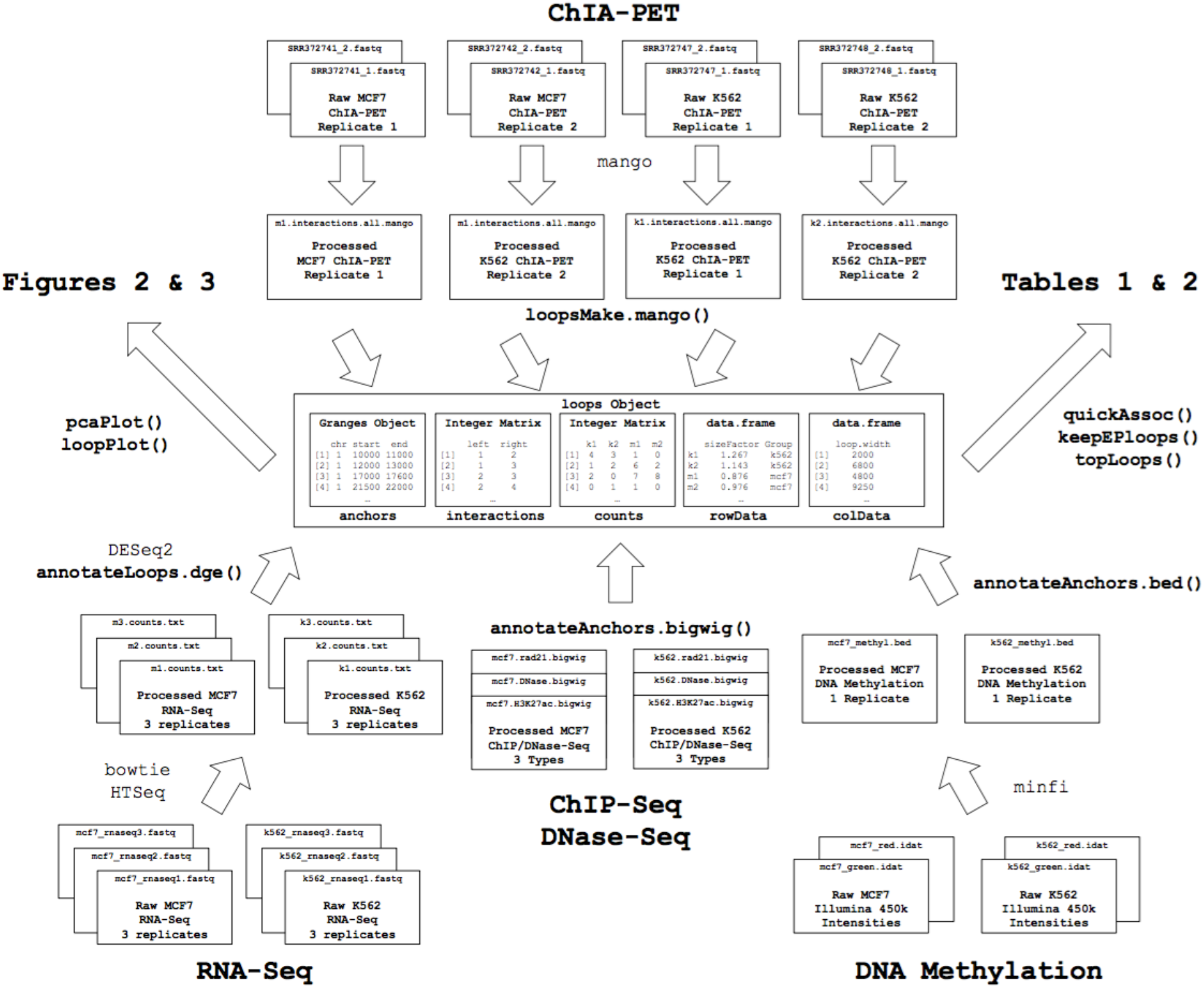
Overview of software, files, data objects, and functions used in the analyses described in this manuscript. Critical functions called in *diffloop* are indicated in bold. External software used in this manuscript to preprocess additional data are also noted in regular font.

### Preprocessing ChIA-PET Sequencing Data for Import into diffloop

A number of software solutions, including the Mango pipeline, exist to process raw sequencing reads from a ChIA-PET assay into putative loops in the bedpe format that is imported into *diffloop* [28]. These tools typically consider interactions for a single sample at a time when identifying statistically significant loops. For the purposes of differential analysis, it is important to make this determination using all samples collectively on order to define a common set of loops. We therefore recommend the use of inclusive preprocessing parameters that do not filter out interactions, retaining all data for import and subsequent filtering in diffloop. In Mango, this is achieved by setting the reportallpairs flag to TRUE.

### Encode ChIA-Pet Data

All ChIA-PET data in this study was generated as part of the ENCODE Project and downloaded from the Sequence Read Archive (SRA)[9]. The format of raw ChIA-PET data is .fastq files that correspond to paired-end reads from a sequencing experiment. For our preprocessing, we used the default parameters in Mango except for specifying that all interactions be preserved (reportallpairs = TRUE) and ChIP peaks be extended by 1,000 base pairs (peakslop = 1000) rather than the default 500 bp. Additionally, we specified linker sequences previously described in the ENCODE ChIA-PET protocol[9] which also correspond to the default parameters in Mango. **Table 4 (A)** provides an overview of the ChIA-PET samples used in this study, including the raw read counts, the location of the data on GEO, as well as the number of PETs used in *diffloop* after data processing with Mango. These results of the Mango processing are also summarized in **Figure 2 (D)**.

**Table 4:**
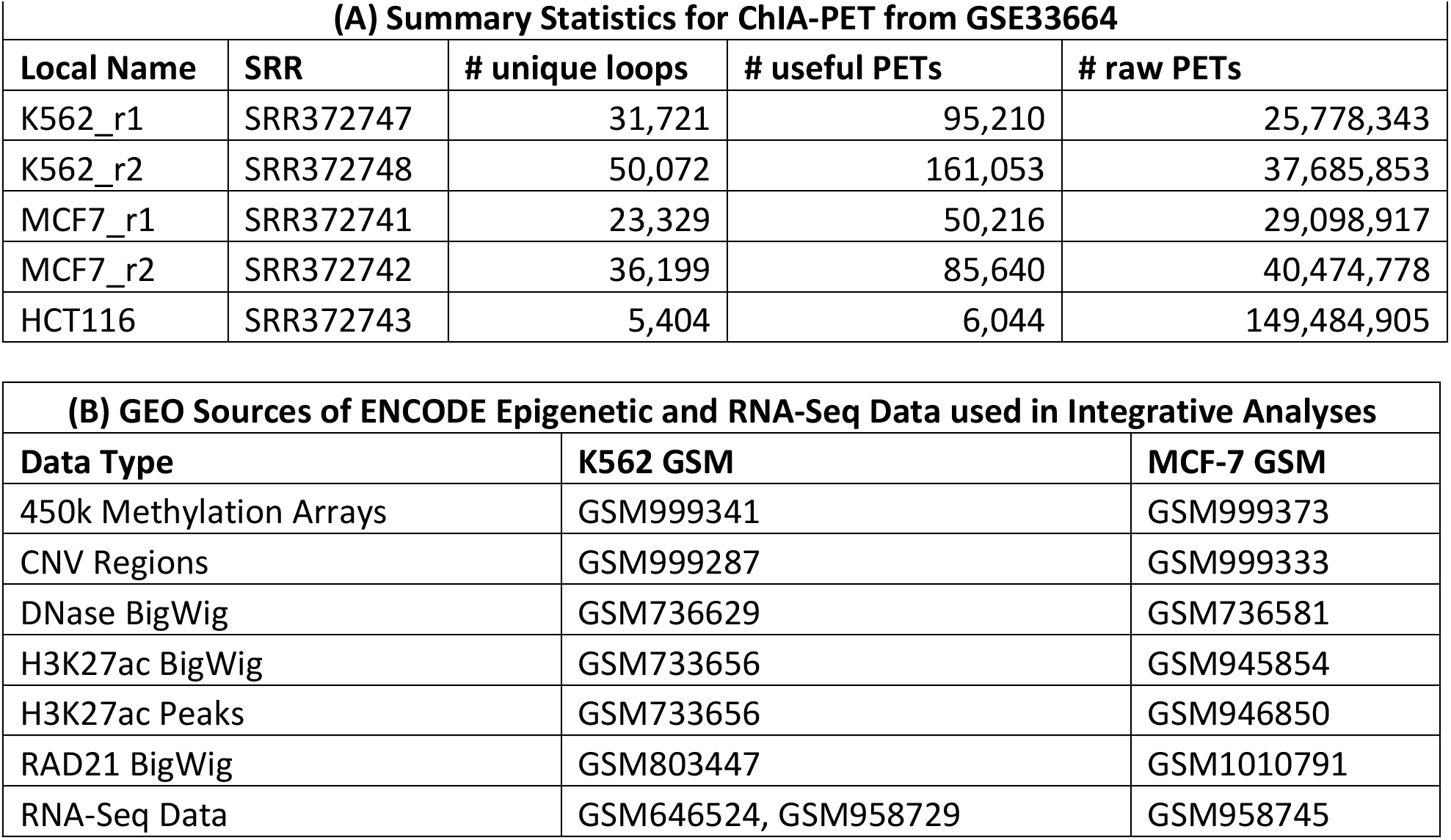
Published data used in this manuscript. (A) Accession of GEO sequencing reads for each of the ChIA-PET samples with summary statistics. (B) Accessions of the epigenetic, genetic, and transcriptomic data used for downstream analyses in *diffloop*.

### Quality Control in diffloop

We filtered amplified or deleted copy number variation (CNV) regions in either of K562 or MCF-7 using the **removeRegion()** function to reduce the chance that genome alterations would bias differential loop calls. We next retained only those loops whose interaction counts were significantly higher than that expected based on the background chromatin interaction frequency using a threshold of 0.01 on q-values generated by the **mangoCorrection()** function. *diffloop* aggregates count data across all samples, providing more power to call valid loops than when analyzing each sample individually. In order to further reduce the multiple testing burden we further restricted loops to those with a minimum of 2 samples with at least 2 PETs per loop (similar as to what was used previously [13]) using the **filterLoops()** function. The low counts associated with the discarded loops would preclude meaningful inference about between group differences.

One peculiar feature of the data was the presence of “discordant” loops that were highly variable between the replicates. Setting a threshold of five or more counts in one replicate but zero in the other identified 337 such loops (red dots on **Figures S3 (A, B)**). Some of these discordant loops appear significantly differential as a result of the variance shrinkage performed in the association model. For example, while a loop with counts of 45 and 0 for one group and 0 and 0 for another is classified as differential, this finding is unlikely to be reliable. Many (166 of 337 or 49%) of these identified discordant loops (such as the example noted above) were removed using the **filterLoops()** function since they don’t meet the criterion of being present in 2+ samples.

### Loop Annotation

The **annotateLoops()** function classifies each loop as enhancer-promoter, promoter-promoter, enhancer-enhancer, or no special annotation for POL2 loops. Additional annotation for loops mediated by CTCF or cohesin subunits is also currently supported. Moreover, loops that connect distal regulatory elements and gene promoters can be selected using the **keepEPloops()** function. For the analyses described in this manuscript, enhancer regions were defined by a 1kb radius around an H3K27ac peak for either cell type. Other epigenetic markings may also be suitable for nuanced logic in defining promoter and regulatory regions. For our analyses, peaks were downloaded from GEO using the accession numbers provided in **Table 4(B)**. Promoter regions were defined as being within a 1kb radius of a RefSeq transcription start site.

### Differential Loop Calling

To determine differential topological features, *diffloop* uses the *edgeR* package to model PET counts from the loops object counts matrix using a negative binomal distribution [14]. To account for variation in sequencing depth, a per-sample size factor is included as an offset in the model, as previously described. Loop-wise dispersion estimates are stabilized using an empirical Bayes shrinkage procedure. In effect, the implementation of differential loop calling in *diffloop* applies the RNA-Seq gene read count test from edgeR to loop PET counts.

Though the application of the edgeR framework to our counts data was intuitive, we note the mean-variance trend in **Figure S1** that fails to show overdispersion beyond that in the Poisson model at the current sequencing depths. As this pattern is also observable at low counts for many RNA-Seq datasets, [29] we expect that topology libraries with better sequencing depths will likely also demonstrate greater overdispersion and benefit from application of the Negative Binomial regression model.

To assess whether alternate models may be better candidates for identifying differential loops, we considered a stratified model using binned loop widths (**Figure S4, S5**) and examined loops with altered significance compared to the standard model (**Table S1, S2**). We also considered the limma-voom model (**Figure S6, S7**) and noted the loops with the largest changes in significance (**Table S3, S4**). We found no evidence that the alternate models performed better than the edgeR model. The supplement provides a more complete synopsis of these analyses.

### Integration of DNA Methylation, DNase Hypersensitivity, and ChIP-Seq Data

*diffloop* provides means for integration of processed epigenetic data as shown in **Figure 6**. To demonstrate this functionality, raw probe intensities from the Illumina 450k array were processed using minfi v1.3.1[30] and exported as .bedgraph format files for both the K562 and MCF-7 cell lines. Per-anchor methylation was computed by averaging the Beta methylation estimates across all CpGs contained in the specific anchor using the **annotateAnchors.bed()** function. Bigwigs of RAD21, H3K27ac, and DNase hypersensitivity were downloaded from GEO accessions as specified in **Table 4 (B)**. Similar to the methylation values, per-anchor intensities for the ChIP-Seq and chromatin accessibility were computed by averaging over all values contained in the specific anchor using the **annotateAnchors.bigwig()** function.

### Differential Expression Analyses

Paired 75 base pair reads from PolyA RNA-Seq for each of the K562 and MCF-7 cell lines from the ENCODE Project were processed (GEO series GSE33480). These data included two samples for K562 (GSM958729) and three samples for MCF-7 (GSM958745). An additional replicate was processed for K562 (GSM646524) for a balanced differential expression analysis. Each samples’ reads were individually aligned using Tophat v1.0.14 [31] and hg19 RefSeq reference transcriptome counts were generated using HTSeq 0.6.1.[32] Differential expression was performed using DESeq2 v1.11.45 [10] to determine variable gene expression between the two cancer cell lines. Enhancer-promoter loops that uniquely linked to a single transcription start site were annotated with the summary statistics from DESeq2 using the **annotateLoops.dge()** function. While this function has additional parameters to handle loops that do not clearly link to a single transcription start site, all analyses including transcription annotation retained only enhancer-promoter loops where the “promoter” anchor mapped to within 1kb of a single transcription start site in the hg19 Refseq build.

## DECLARATIONS

### List of abbreviations

- 3C: Chromatin Conformation Capture
- 3D: Three-dimensional
- bedpe: BED paired-end (file format)
- ChIA-PET: Chromatin Interaction Analysis by Paired-End Tag Sequencing
- ChIP: Chromatin Immunoprecipitation
- ChIP-seq: Chromatin Immunoprecipitation Sequencing
- CPT: ChIA-PET Tool
- CNV: Copy Number Variation
- FDR: False Discovery Rate
- GEO: Gene Expression Omnibus
- Hi-C: high-throughput chromosome conformation capture
- kb: kilobase
- PC: Principal Components
- PCA: Principal Component Analysis
- PET: Paired-end tag (paired-end sequencing read)
- RNA Pol II: RNA Polymerase II
- Seq: Sequencing
- SRA: Sequence Read Archive
- TAD: Topologically Associated Domain

### Ethics Declaration

Not applicable

### Consent for publication

Not applicable

### Availability of data and material

*diffloop*, is available from R/Bioconductor: https://bioconductor.org/packages/release/bioc/html/diffloop.html. This package requires R > = 3.3.0 and depends on several R/Bioconductor packages including GenomicRanges, rtracklayer, and ggplot2. Version 1.3.2 of *diffloop* was used for all analyses described in this manuscript. All data used in this study are made freely available through GEO at the accession numbers specified in the Methods section. Code to reproduce analyses, figures, and tables are available through Github: **https://github.com/aryeelab/diffloop_paper/**

## Author contributions

CAL and MJA conceived and designed the study. CAL and MJA wrote the software, analyzed data, and interpreted the results. CAL and MJA both wrote the manuscript.

## Acknowledgements

We thank D. Day, D. Hnisz, and members of the R. Young Lab for their useful discussion and insight.

## Additional Files

**Additional File 1. PDF. *diffloop* Supplement**. Supplementary information regarding additional model diagnostics considered in the differential loop calling implementation.

## References

1. Matharu N, Ahituv N: Minor Loops in Major Folds: Enhancer-Promoter Looping, Chromatin Restructuring, and Their Association with Transcriptional Regulation and Disease. PLoS Genet 2015, 11:e1005640.

2. Flavahan WA, Drier Y, Liau BB, Gillespie SM, Venteicher AS, Stemmer-Rachamimov AO, Suva ML, Bernstein BE: Insulator dysfunction and oncogene activation in IDH mutant gliomas. Nature 2016, 529:110–114.

3. Wang S, Wen F, Wiley GB, Kinter MT, Gaffney PM: An enhancer element harboring variants associated with systemic lupus erythematosus engages the TNFAIP3 promoter to influence A20 expression. PLoS Genet 2013, 9:e1003750.

4. Hnisz D, Weintraub AS, Day DS, Valton AL, Bak RO, Li CH, Goldmann J, Lajoie BR, Fan ZP, Sigova AA, et al: Activation of proto-oncogenes by disruption of chromosome neighborhoods. Science 2016, 351:1454–1458.

5. Dekker J, Rippe K, Dekker M, Kleckner N: Capturing chromosome conformation. Science 2002, 295:1306–1311.

6. Rao SS, Huntley MH, Durand NC, Stamenova EK, Bochkov ID, Robinson JT, Sanborn AL, Machol I, Omer AD, Lander ES, Aiden EL: A 3D map of the human genome at kilobase resolution reveals principles of chromatin looping. Cell 2014, 159:1665–1680.

7. Durand NC, Robinson JT, Shamim MS, Machol I, Mesirov JP, Lander ES, Aiden EL: Juicebox Provides a Visualization System for Hi-C Contact Maps with Unlimited Zoom. Cell Syst 2016, 3:99–101.

8. Huber W, Carey VJ, Gentleman R, Anders S, Carlson M, Carvalho BS, Bravo HC, Davis S, Gatto L, Girke T, et al: Orchestrating high-throughput genomic analysis with Bioconductor. Nat Methods 2015, 12:115–121.

9. Consortium EP: An integrated encyclopedia of DNA elements in the human genome. Nature 2012, 489:57–74.

10. Love MI, Huber W, Anders S: Moderated estimation of fold change and dispersion for RNA-seq data with DESeq2. Genome Biol 2014, 15:550.

11. Wu HJ, Michor F: A computational strategy to adjust for copy number in tumor Hi-C data. Bioinformatics 2016.

12. Phanstiel DH, Boyle AP, Heidari N, Snyder MP: Mango: a bias-correcting ChIA-PET analysis pipeline. Bioinformatics 2015, 31:3092–3098.

13. Ji X, Dadon DB, Powell BE, Fan ZP, Borges-Rivera D, Shachar S, Weintraub AS, Hnisz D, Pegoraro G, Lee TI, et al: 3D Chromosome Regulatory Landscape of Human Pluripotent Cells. Cell Stem Cell 2016, 18:262–275.

14. Robinson MD, McCarthy DJ, Smyth GK: edgeR: a Bioconductor package for differential expression analysis of digital gene expression data. Bioinformatics 2010, 26:139–140.

15. Law CW, Chen Y, Shi W, Smyth GK: voom: Precision weights unlock linear model analysis tools for RNA-seq read counts. Genome Biol 2014, 15:R29.

16. Subramanian A, Tamayo P, Mootha VK, Mukherjee S, Ebert BL, Gillette MA, Paulovich A, Pomeroy SL, Golub TR, Lander ES, Mesirov JP: Gene set enrichment analysis: a knowledge-based approach for interpreting genome-wide expression profiles. Proc Natl Acad Sci U S A 2005, 102:15545–15550.

17. Sun J, Nawaz Z, Slingerland JM: Long-range activation of GREB1 by estrogen receptor via three distal consensus estrogen-responsive elements in breast cancer cells. Mol Endocrinol 2007, 21:2651–2662.

18. Sengupta S, Sharma CG, Jordan VC: Estrogen regulation of X-box binding protein-1 and its role in estrogen induced growth of breast and endometrial cancer cells. Horm Mol Biol Clin Investig 2010, 2:235–243.

19. Delgado MD, Leon J: Myc roles in hematopoiesis and leukemia. Genes Cancer 2010, 1:605–616.

20. Moon HW, Kim TY, Oh BR, Min HC, Cho HI, Bang SM, Lee JH, Yoon SS, Lee DS: MTHFR 677CC/1298CC genotypes are highly associated with chronic myelogenous leukemia: a case-control study in Korea. Leuk Res 2007, 31:1213–1217.

21. Rossen RD, Laughter AH, Orson FM, Flagge FP, Cashaw JL, Sumaya CV: Human peripheral blood monocytes release a 30,000 dalton factor (30 KD MF) that stimulates immunoglobulin production by activated B cells. J Immunol 1985, 135:3289–3297.

22. Curran JE, Weinstein SR, Griffiths LR: Polymorphic variants of NFKB1 and its inhibitory protein NFKBIA, and their involvement in sporadic breast cancer. Cancer Lett 2002, 188:103–107.

23. Bell JT, Pai AA, Pickrell JK, Gaffney DJ, Pique-Regi R, Degner JF, Gilad Y, Pritchard JK: DNA methylation patterns associate with genetic and gene expression variation in HapMap cell lines. Genome Biol 2011, 12:R10.

24. Li G, Ruan X, Auerbach RK, Sandhu KS, Zheng M, Wang P, Poh HM, Goh Y, Lim J, Zhang J, et al: Extensive promoter-centered chromatin interactions provide a topological basis for transcription regulation. Cell 2012, 148:84–98.

25. Lun AT, Smyth GK: diffHic: a Bioconductor package to detect differential genomic interactions in Hi-C data. BMC Bioinformatics 2015, 16:258.

26. Aran D, Sabato S, Hellman A: DNA methylation of distal regulatory sites characterizes dysregulation of cancer genes. Genome Biol 2013, 14:R21.

27. Lawrence M, Huber W, Pages H, Aboyoun P, Carlson M, Gentleman R, Morgan MT, Carey VJ: Software for computing and annotating genomic ranges. PLoS Comput Biol 2013, 9:e1003118.

28. Li G, Fullwood MJ, Xu H, Mulawadi FH, Velkov S, Vega V, Ariyaratne PN, Mohamed YB, Ooi HS, Tennakoon C, et al: ChIA-PET tool for comprehensive chromatin interaction analysis with paired-end tag sequencing. Genome Biol 2010, 11:R22.

29. Anders S, Huber W: Differential expression analysis for sequence count data. Genome Biol 2010, 11:R106.

30. Aryee MJ, Jaffe AE, Corrada-Bravo H, Ladd-Acosta C, Feinberg AP, Hansen KD, Irizarry RA: Minfi: a flexible and comprehensive Bioconductor package for the analysis of Infinium DNA methylation microarrays. Bioinformatics 2014, 30:1363–1369.

31. Trapnell C, Pachter L, Salzberg SL: TopHat: discovering splice junctions with RNA-Seq. Bioinformatics 2009, 25:1105–1111.

32. Anders S, Pyl PT, Huber W: HTSeq—a Python framework to work with high-throughput sequencing data. Bioinformatics 2015, 31:166–169.

